# Accelerating Prediction of Chemical Shift of Protein Structures on GPUs

**DOI:** 10.1101/2020.01.12.903468

**Authors:** Eric Wright, Mauricio Ferrato, Alex Bryer, Robert Searles, Juan Perilla, Sunita Chandrasekaran

**Affiliations:** Dept. of CIS, UDEL, Newark, DE, USA; Dept of Chemistry, UDEL, Newark, DE, USA; NVIDIA, CA, USA

## Abstract

Experimental chemical shifts (CS) from solution and solid state magic-angle-spinning nuclear magnetic resonance spectra provide atomic level information for each amino acid within a protein or protein complex. However, structure determination of large complexes and assemblies based on NMR data alone remains challenging due the complexity of the calculations. Here, we present a hardware accelerated strategy for the estimation of NMR chemical-shifts of large macromolecular complexes based on the previously published PPM_One software. The original code was not viable for computing large complexes, with our largest dataset taking approximately 14 hours to complete. Our results show that the code refactoring and acceleration brought down the time taken of the software running on an NVIDIA V100 GPU to 46.71 seconds for our largest dataset of 11.3M atoms. We use OpenACC, a directive-based programming model for porting the application to a heterogeneous system consisting of x86 processors and NVIDIA GPUs. Finally, we demonstrate the feasibility of our approach in systems of increasing complexity ranging from 100K to 11.3M atoms.

**Author summary:** 

## Introduction

Computing architectures are ever-evolving. As these architectures become increasingly complex, we need better software stacks that will help us seamlessly port real-world scientific applications to these emerging architectures. It is also important to prepare applications such that they can be readily retargeted to existing and future systems without the need for drastic changes to the code itself. In an ideal world, we are looking for solutions to create a performance productive software. However, this is not easy and is sometimes an impossible task to accomplish.

Programming and optimizing for different architectures at a minimum often require codes written in different programming languages. This presents an inherent difficulty for software developers as they would need to develop and maintain an entire secondary code base. For this reason, it is ideal to have a single programming standard that is both portable to all architectures and maintains high performance. There are three main reasons why this is difficult: (1) Sufficient parallelism is not exposed to hardware architecture if the algorithm is structured in a way that it limits the level of concurrency, (2) Features in a programming model are often hardware-facing and only occasionally application/user-facing, and (3) to encompass different applications from different fields of study would require the programming standard to have many levels of abstraction in a sensible way.

There are currently three widely accepted solutions that software developers adapt create performance portable applications: libraries, languages, and directives. Libraries suffer from an inherent scope problem; they can only solve a specific subset of problems and are only designed for a specific subset of architectures. Languages are flawed because of the reasons previously outlined such as requiring the programmer to rewrite significant amounts of code. Directives are special lines of code added alongside standard code. These additional lines of code act as hints to the compiler that create the necessary executables for the platform the code is being compiled upon. This provides the portability that is important to software developers and in many applications offer competitive performance when compared to its language-specific counterpart. We believe that directives offer a reliable balance between performance and portability and is what we will focus on for the remainder of this manuscript.

Two directive-based programming models that are widely popular are OpenMP [1, 2] and OpenACC [3]. OpenMP was created in 1997 as a shared-memory programming model. Since 2013 (OpenMP 4.0 offloading), the model has begun to target heterogeneous computing systems and is continuing to evolve. Applications that have been deployed using the OpenMP offloading model include Pseudo-Spectral Direct Numerical Simulation-Combined Compact Difference (PSDNS-CCD3D) [4] - a CFD code for turbulent flow simulation, and Quicksilver [5] - a Monte Carlo Transport code. OpenACC is the other directive-based programming model and was created in 2011. The model has since been adopted widely by scientific developers, to port their large scientific applications—sometimes production code—to heterogeneous architectures. Some examples include ANSYS [6], GAUSSIAN [7], nuclear reactor code Minisweep [8] (mini application of Denovo), and Icosahedral non-hydrostatic (ICON) [9]. Both OpenMP and OpenACC allow incremental improvement to a given code base. Directives also help create a re-usable code especially when the implementations can target more than one type of architecture.

A few points to note: For the current manuscript, we have chosen the OpenACC model after observing the OpenACC compiler’s (PGI implementation) maturity and stability. GCC (by Mentor Graphics) also offers an OpenACC implementation, however at the time of running this experiment the implementation was not yet mature enough.

### Overview of the Scientific Problem: Chemical Shift Prediction

Nuclear magnetic resonance (NMR) is an experimental technique employed in numerous fields such as chemistry, physics, biochemistry, biophysics and structural biology. A chemical shift, the principle observable in NMR instrumentation, provides valuable insight into protein secondary structure by allowing inference about conformation to be drawn based on peak shift. Measured in parts per million (ppm), a chemical shift describes the resonant frequency of a nucleus by comparing its observed frequency to that of a standard reference in the presence of a magnetic field. Magnetic resonance imaging, or MRI, is a familiar application of this powerful technology.

Computational tools to aid structure determination with NMR observables have materialized into a rich domain of protein study and protein chemical shifts have been used in varying ways to successfully elucidate structure. Commonly, these programs employ perusal of scientific databases to establish and parse relationships between shifts, sequence and structure [10–15]. The accuracy of such solutions is contingent upon the availability of data, both from NMR experiment and sequence homology assignment, the latter which treats the similarity among proteins according to commonalities in their sequences and probable evolution. Thanks to projects such as the BioMagResBank (BMRB) [16], NMR data is more available than ever before, engendering the feasibility of semi-empirical prediction methods which utilize existing chemical shift data to parameterize functional prediction models.

Obviating the need for database searching and sequence matching is a semi-empirical method named PPM [17]. The goal of PPM was to provide a prediction model that could operate over NMR conformational ensembles, predict chemical shifts from structure and provide new dimensions of protein forcefield refinement, structural refinement, and ensemble validation—a goal which PPM met aptly. In a departure from ensemble analysis, PPM’s successor PPM_One introduced a static-structure based chemical shift prediction method that showed competitive accuracy with other software [18].

### Motivation

Drawing from approximations of first principle calculations and trained with accessible NMR data, the PPM_One model considers chemical shift as a sum of discrete *descriptors*. These descriptors, which quantify chemical shifts due to ring current effects, hydrogen bond effects, dihedral angles, and more [17, 18], take the form of relatively simple— and differentiable—functions of the atomic coordinates. Considering these factors, PPM_One is a prime target for parallelization and optimization; to extend practical application of the software to larger structures, populous NMR ensembles, or molecular dynamics trajectories describing thousands of structures. While a suitable candidate to this end, the original PPM_One code was not written in a way to exploit the massive compute power of accelerators such as GPUs. In our work, we have ported the PPM_One application to utilize parallel hardware, such as GPUs, using OpenACC.

### Contributions

- Equip domain scientists with an accelerated version of PPM_One that functions in a realistic lab environment.
- Provide an accelerated chemical shift prediction code that can be adapted to large Molecular Dynamics packages.
- Demonstrate the feasibility and scalability of our approach in systems of increasing complexity ranging from 2,000 to 13,000,000 atoms.

## Materials and methods

### Preparing code for acceleration

Before accelerating or parallelizing a given code, a standard practice is to identify computational hotspots that take the most execution time. This generally means that we are looking for the largest or most intensive loops that exist within the application. To find these portions of the code we use dedicated profiling tools, then we examine the source code and determine if they are or could be refactored to be accelerator-friendly.

### Identifying Computational Hotspot in Chemical Shift Prediction

The OpenACC-enabled profiler that comes packaged with the PGI compiler is called PGPROF. PGPROF displays detailed information about CPU and GPU performance. This information includes breakdowns by runtime, memory management, and accelerator utilization. We used PGPROF to find functions in PPM_One that are the most time consuming and thus are the most important parallelization targets. The two main functions (1) *predict bb static ann*() and (2) *predict proton static new*() accounted for the majority of the total runtime (81.23% and 16.28% respectively). These two functions are composed of other smaller functions that were also analyzed using PGPROF. The most significant of these was *get_contact*(), taking 35% of the total runtime. We also observed that the time taken for *get_contact*() scales well with the dataset size. When profiling with large molecules (1+ million atoms), we found that *get_contact*() could take upwards of 80% of the programs total runtime. This makes *get_contact*() the most important sub-function and our first target for parallelization. *get select*() follows with 23% of runtime, but it was optimized by our initial code refactoring and will be outlined in Section 2.2.

Fig 1 shows the results of our profile when using a relatively small molecule (100,000 atoms). Some other functions that were found in the profile are *getani*() (18% runtime), *gethbond*() (15% runtime), and *getring*() (12% runtime). Similar to *get_contact*(), we also found that *gethbond*() scales somewhat well with the size of the molecule, whereas *getani*() and *getring*() do not scale as much.

**Fig 1.**
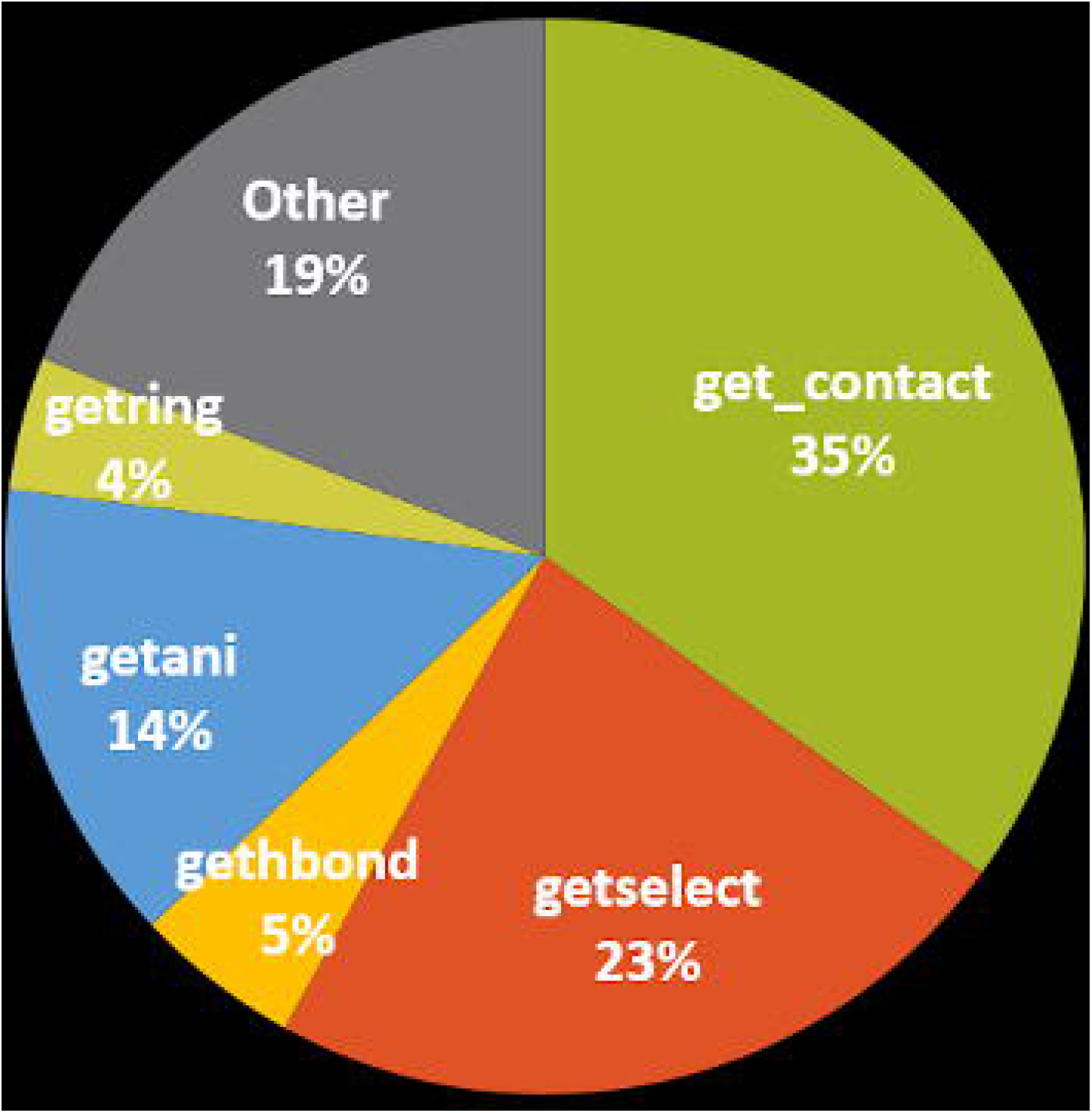
Visual representation of function runtime breakdown before and after serial optimizations.

### Inital Code Refactoring

Many of these functions in the original sequential code were written as a direct implementation of their respective algorithms. Unfortunately, this caused issues when attempting to move the code to accelerators. One of our first observations when using the profiling tools was a significant redundancy of memory copying caused by calling the *getselect*() function an unnecessary number of times. To fix this, we altered the code to only call *getselect*() once, and then store and reuse the associated memory. This optimization alone led to a 20% performance increase when running with some of the datasets.

The next optimization we made was to a function called *clear*(). The *clear*() function filters through a list of protons, removing any of them that do not work with the algorithm. The way that the protons were removed from the list was simply inefficient; the runtime of this function varied greatly depending on which dataset was tested since some molecular structures require more protons to be filtered than others. The *clear*() function could vary from taking seconds to taking hours depending on the dataset alone. As a result, we rewrote *clear*() to use a more efficient list filter that made the operation take only a few seconds or less for all structures.

Lastly, we ran into some problems with the C++ STL containers that were used within the code. This mostly applied to the C++ standard vector class. To account for this, many C++ vectors were replaced with basic arrays. This did not have any meaningful impact on the performance on its own but allowed for more efficient communication with the GPU. In other places, we interfaced with the vector containers by using the built-in *data*() function to retrieve the underlying memory, allowing us to move the data to the GPU without the need to use extra libraries or code rewrites.

### Using OpenACC for Acceleration

This section begins with a brief overview of the OpenACC model followed by using OpenACC for acceleration.

### Overview of OpenACC model

As mentioned earlier, OpenACC is a directive-based programming model that targets heterogeneous systems comprising of CPUs and accelerators. The model exposes three levels of parallelism via gang, worker and vector parallelism that enables programmers to abstract the architecture along with maximally utilizing the potential of multicore or accelerators. Since the model is directive-based it allows the programmers to achieve performance while almost maintaining the original source code base. Typically, compute-intensive portions of the program often identified by profilers are offloaded to the accelerators; a task orchestrated by the host by allocating memory on the accelerator device, initiating data transfer, offloading the code to the accelerator, passing arguments to the compute region, queuing the device code, waiting for completion, transferring results back to the host, and deallocating memory. With often only minor adjustments to memory management near parallelized compute regions, the model accommodates both shared and discrete memory or any combination of the two across any number of devices. The model has the capacity to expose the separate memories through the use of a device data environment.

### Acceleration

The following section highlights the usage of some of the OpenACC features for our case. They are also the most commonly used features.

After ensuring that the code was accelerator-friendly, we began applying OpenACC directives to the code. We tackled each function individually in order of importance, meaning that we started with *get_contact*() and finished with *getring*(). Everytime we made a meaningful alteration we would re-run the code on a few different datasets and compare the results to their non-accelerated baselines. This would let us know if we made any errors along the way.

We decorated the major loops in the code with the OpenACC *parallel loop* directive. This told the compiler to offload these loops on the GPU automatically. In some cases, this alone was enough to see a speedup as some loops were embarassingly parallel. However, in other cases we saw significant slowdown and sometimes wildly incorrect code output compared to our serial baseline. These two problems were overcome by using other OpenACC features.

To fix our incorrect output, we had to implement both the *reduction clause* and *atomic directive*. The reduction clause is important to include in parallel loops that contain race conditions. These are areas in the code that can result in errors when multiple parallel units overwrite each other in shared memory. The reduction clause prevents this by aligning memory reads/writes to produce a single coherent value.

The atomic directive fills a similar purpose. However, it is useful in situations where many different race conditions could occur at different locations in memory. There was only one situation in our code where a reduction clause was not sufficient, and that was in the *gethbond*() function.

The other problem we overcame was handling overall slowdown in the code. This is largely due to having too many memory transfers between the host and device. After profiling our initial parallelization of the *get_contact*() function, we saw that the majority of the time was spent on transferring data between the host and device memory. Originally, *get_contact*() would be called many times throughout code execution (hundreds to thousands of times, depending on the dataset). We added a loop that would iterate over all of the individual *get_contact*() calls, which gave us another dimension to expose parallelism. This also means that no data would need to be transferred between the different calls of *get_contact*(). This change was beneficial because out of all of the functions *get_contact*() received the largest speed-up. The speed-up will be discussed in more detail in Section 3.2.

To further elaborate on our memory management, we originally started with a simple strategy; copy everything to the device that was needed immediately before the loop starts, and then copy everything back to the CPU that will be needed elsewhere. This strategy proved to perform badly as much of the data needed on the device was being moved multiple times unnecessarily. We changed this to instead transfer the data after the code’s preprocessing; before any of the main computation happens. Then, we transfer the computation results results back to the host so that it can then be printed to files.

### Target Functions for Acceleration

Each of the functions we have identified are important to the overall chemical shift prediction algorithm that PPM_One implements. *get_contact*() is one of the most important functions in the PPM_One algorithm due to the fact that it serves as the principle interface between the input coordinates and secondary structure contact data. *get_contact*() iterates over all atomic positions, given in the molecule, and computes a distance between each atom index and the successive atom index. Next, for each atom in each residue in the PPM_One input structure, the random-coil chemical shift for atoms in that residue is applied as a fit parameter to normalize the calculated chemical shift, ultimately ascertaining local flexibility given that more disordered structures (or regions of the structure) will have chemical shifts tending towards the random-coil chemical shift value. Since this procedure must be carried out exhaustively over the entire structure and manages data from individual function calls and parameter tables, it takes up a proportionally large piece of the total runtime. Adequate parallelization of this function is of high importance as otherwise it poses a large sequential-bottleneck in the total runtime of the program.

*gethbond*() computes the effect that backbone hydrogen bonding has on chemical shift. PPM_One describes this effect in terms of the inverse of donor-acceptor distance, and applies a descriptor based on the angle formed between two different atom triples, NHO and *HOC′*. Since every amino acid has donor-acceptor pairs, this function gets called with high frequency and involves distance and angle calculations for each donor and acceptor relative to the specified atom triples, making *gethbond*() a meaningful target for parallelization and performance-gain despite its relatively simple formulation.

The function *getani*() represents the compute region for calculating the chemical shift due to magnetic anisotropy. Magnetic anisotropy quantifies the directionally-dependent electromagnetic interactions between atoms. PPM_One employs this calculation for interactions between protons and peptide-amide groups consisting of Oxygen (*O*), Carbon prime (*C′*) and Nitrogen (*N*). Additional calls are made to *getani*() for side-chain *OCN* groups of Asparagine and Glutamine, *OCO* side-chain groups of Glutamate and Aspartate, and the *NCN* side-chain of Arginine. The formulation for the calculation used by PPM_One is known as the “axially symmetric model” [19], in accordance with McConnell’s characterization of anisotropy of peptide groups [20]. At each function call, the distance between the queried proton and the peptide-amide group is calculated. This, the vectors pointing from the proton to the peptide-amide group, and from the proton to the normal vector of the peptide-amide are used to compute an angle to pass into the magnetic anisotropy expression.

*getring*() encompasses two different functions in the PPM_One program that calculate the chemical shift due to ring-current effects; one function calculates ring-current effects felt by Hydrogen atoms with respect to an aromatic residue, and the other calculates the effect felt by backbone atoms adjacent to an aromatic residue. PPM_One considers the aromaticity of amino acids Phe, Tyr, His, Trp-5 and Trp-6. The aromatic rings of these residues have important structural implications due to electrostatic induction, as the circular movement of delocalized electrons (ie, current) in conjugated Pi-bonding orbitals induces a magnetic field vector orthogonal to the plane described by the atoms of the ring. To quantify this effect, the queried atom’s position in cartesian space must be projected to a position on the 2D subspace defined by the plane of the aromatic ring. Additionally, distances between all atoms in the ring are calculated in this function each time it is called, making it costly to compute even though its application is limited to only aromatic residues and atoms in their local environment.

## Results

This section will elaborate on the experimental setup and the results obtained.

### Experimental Setup

For the multicore, V100 and P40 results shown in both the tables, we use the PSG DGX-1b compute node consisting of Intel Xeon e5-2698 v4 20 cores and a single NVIDIA Volta V100 card and another compute node that has a single P40. For the serial runs shown in both the tables, since we could not get time on the PSG system, we have used our internal University of Delaware’s local system that has an Intel x990 core.

### Datasets

Fig 2 shows the different datasets used for our experiments, represented to scale. The first tested dataset constitutes 100,000 atoms, roughly a quarter-turn, of the Dynamin GTPase (structure E) extracted and written to their own Protein Database (PDB) file. The next dataset tested, structure B, was the HIV-1 capsid assembly (CA) without Hydrogens. This structure was tested without Hydrogens for two reasons: 1) to limit the number of atoms for this test case and 2) to create a variety in the swath of tested structures. In regard to the latter, this directly affects the PPM_One algorithm’s treatment of Hydrogen bond effects, tabulated through the *gethbond()* function. Structures C and D correspond to two variants of the HIV-1 CA, Hydrogens included. Structure C is the HIV-1 CA decorated with Cyclophilin A (CypA), structure D is the same HIV-1 CA decorated with Myxovirus resistance protein B (MxB). These two datasets, 5.1 and 5.9 million atoms respectively, were chosen as test cases of heterogenous systems in addition to their increased atom counts compared to the undecorated HIV-1 CA. The HIV-1 CA test-structures are shown next to their dimeric building block 2KOD (structure A), illustrating the ranging scale and complexity of atomistic representations of biomolecules. Finally, the largest two test systems were built from the Dynamin GTPase. Structure E is a 6.8 million atom model, 14 turns, of the GTPase. The largest structure, containing 13.6 million atoms, constitutes 28 turns of the Dynamin GTPase. The secondary-structure of 2KOD was calculated using Stride [21]. All images were rendered using VMD 1.9.4 and the co-distributed, Tachyon parallel ray-tracing library [22, 23].

**Fig 2.**
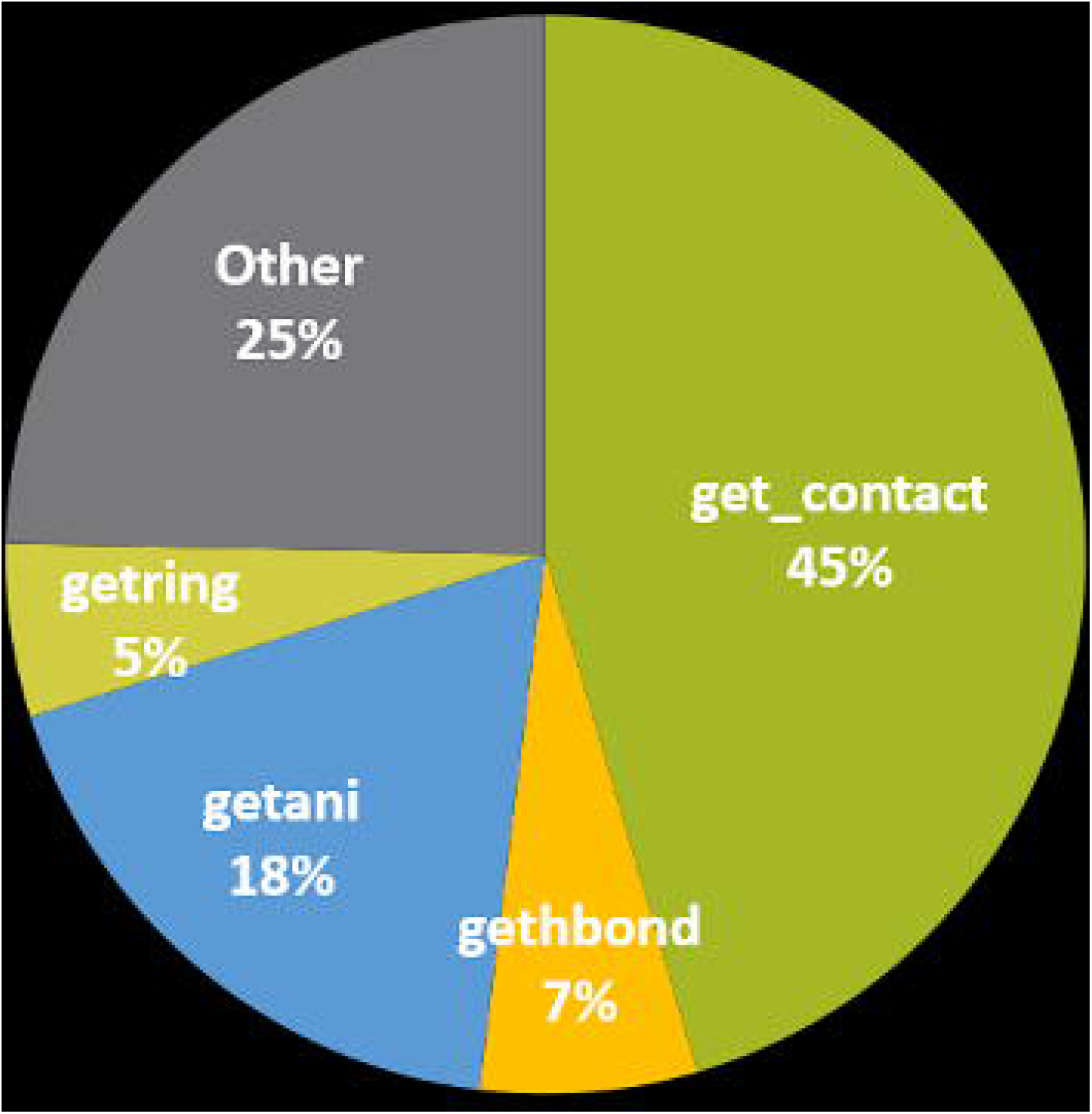
Datasets.

**Fig 3.**
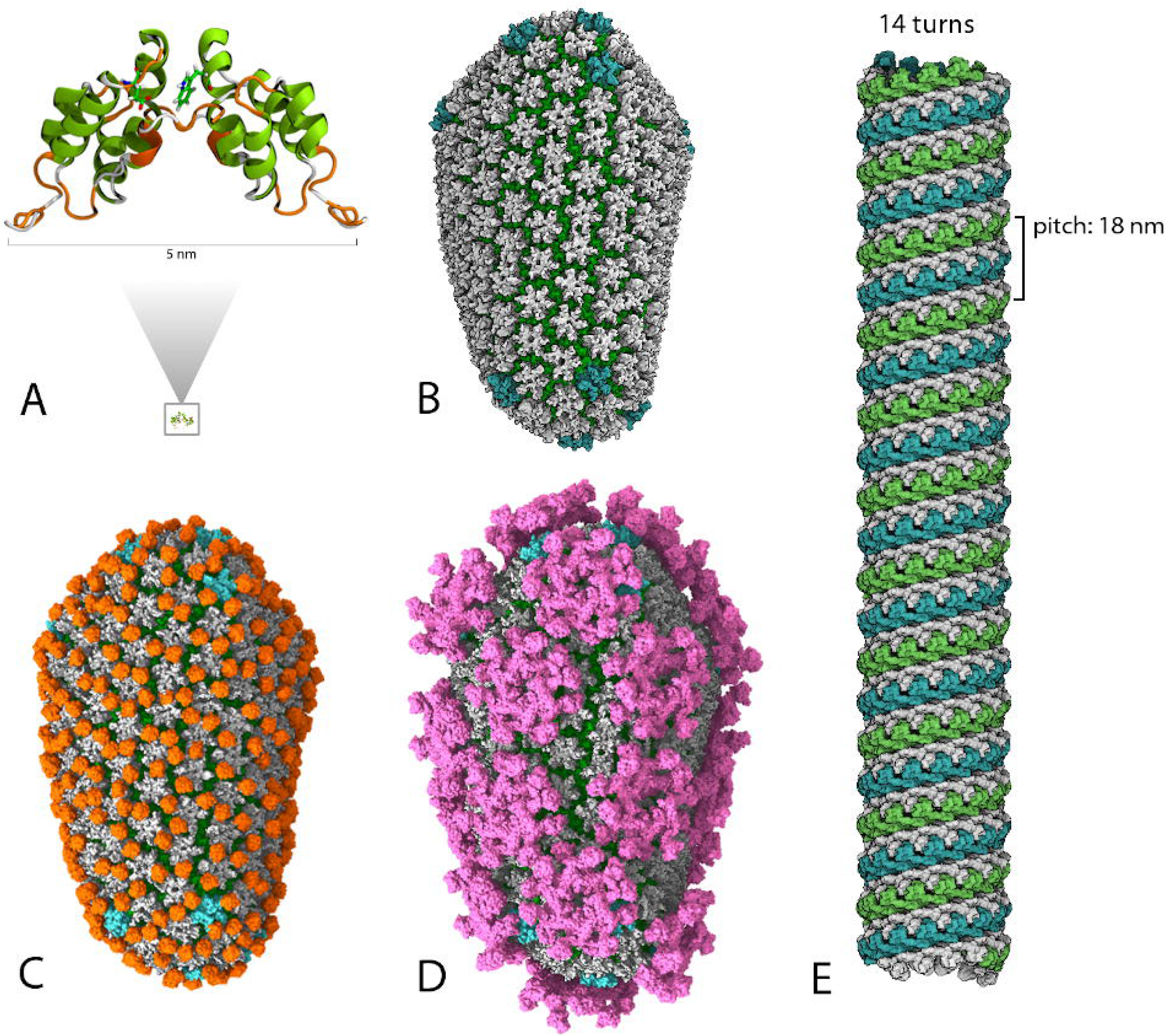
RMSE calculation.

When running the PPM_One application we noticed that the total runtime is proportional to the number of atoms contained in the molecule. However, this is not the only deciding factor. To accommodate for this we are mostly concerned with performance increase of a molecule on different platforms and less concerned with comparing different molecules to each other.

When observing Table 1 we see a significant decrease in total runtime when comparing the serial (optimized) run to any of the accelerators. The multicore performance was 18x faster than the single core results. The Volta V100 results were 56x faster than single core, and 3.1x faster than multicore.

**Table 1.** Results for Small to Large Dataset

|  | 100k atoms | 1.5m atoms | 5m atoms | 6.8m atoms | 11.3m atoms |
| --- | --- | --- | --- | --- | --- |
| Serial (Unoptimized) | 167.11s | 572.01s | 3547.07s | 7 hrs (esimate) | 14 hrs (estimate) |
| Serial (Optimized) | 53.57s | 196.12s | 2003.6s | 1510.71s | 2614.4s |
| Multicore | 4.67s | 32.82s | 116.66s | 153.8s | 146.06s |
| P40 | 3.47s | 17.15s | 56.2s | 78.57s | 72.55s |
| V100 | 3.11s | 13.62s | 39.79s | 49.63s | 46.71s |

When observing individual function performance we see more significant speedup numbers as shown in Table 2. Comparing V100 results to the multicore results, the *get_contact*() function was sped up by 258x, *gethbond*() by 11x, *getani*() by 10x and *getring*() by 3x. Such a high speed up is common for functions that are purely compute intensive and hence can be easily optimized for GPUs. Since our major computational functions are seeing this amount of increase, we predict that much of the remaining total runtime is bound by other portions of the code such as file I/O or preprocessing. We have improved these parts of the code significantly since the start of this project (as seen when comparing the serial unoptimized numbers against the serial optimized). We do not believe that too much more could be done to improve these aspects without rewriting large portions of the code.

**Table 2.** Results for Small Dataset

| 5m atoms | Total Runtime | get_contact | getani | getring | gethbond |
| --- | --- | --- | --- | --- | --- |
| Serial (Optimized) | 2003.60 | 1177.61s | 58.95s | 22.53s | 708.07s |
| Multicore | 116.66s | 51.73s | 2.4s | 0.6s | 25.39s |
| P40 | 56.2s | 1.69s | 1.06s | 0.5s | 17.05s |
| V100 | 39.79s | 0.2s | 0.24s | 0.18s | 2.35s |

### Validation of Results: Calculation RMSE

To calculate the Root Mean Square Error (RMSE), we ran the unaltered code on a single core of a single CPU on 299 different PDB files. Then we reran each file with the developed OpenACC code on the same CPU core, but now with GPU offloading. The following numbers shown in Fig **??** are collected by using the RMSE formula on every prediction of every file comparing the CPU and GPU output.

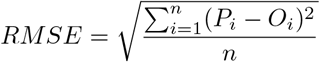

### Source code and Datasets

The PDB files have been previously published and can be found here [24]. The code used for this manuscript is available via our GitHub https://github.com/UD-CRPL/ppm_one.

## Conclusion

PPM_One is a code base that is not written with parallel processors and accelerators in mind. The code base focuses on the chemical shift algorithm. This paper studies the algorithm, profiles the serial code, identifies the hotspots to offload them to multicore and accelerators that make a heterogeneous system. As the model allows the programmer to insert hints to the code, it helps preserve the original code base to a large extent. Such an approach is highly appreciated by domain experts who do not need to learn to nitty-gritties of the architecture before applying such directive-based model, OpenACC.

Scientifically, obtaining these predicted chemical shift values are important for researchers, and if they must wait hours or even days to obtain this information, it can not be used efficiently as a lab utility. Accelerating this program allows users to receive chemical shift information on extremely large data sets. Most importantly, it enables researchers to run these simulations several times every hour, greatly expanding the practical use of this algorithm. The accelerated code can also be called within large molecular dynamics packages allowing the algorithm to be expanded into other codes and applications.

|  | C<sub>α</sub> | C<sub>β</sub> | C' | HN | N | H<sub>α</sub> |
| --- | --- | --- | --- | --- | --- | --- |
| RMS error<br>(ppm) | 1.58e-4 | 8.48e-5 | 1.97e-4 | 5.22e-5 | 2.84e-4 | 1.02e-4 |
| Max error<br>(ppm) | 0.013 | 0.008 | 0.017 | 0.007 | 0.025 | 0.013 |

## Acknowledgments

We gratefully acknowledge the support of NVIDIA PSG Cluster with the donation of the P100 and V100 GPUs used for this research. We are grateful to NVIDIA Corporation for the donation of NVIDIA GPU such as Tesla K40 and a GTX Titan X (Maxwell) GPU cards via the NVIDIA hardware grant program. This material is based upon work supported by the National Science Foundation (NSF) under Grant No. 1814609. The work is also supported by the NIGMS and NIAID (P50GM082251 and P30GM110758). This work used the Extreme Science and Engineering Discovery Environment(XSEDE), which is supported by the National Science Foundation (NSF grant OCI-1053575).

